# Hybridization-based In Situ Sequencing (HybISS): spatial transcriptomic detection in human and mouse brain tissue

**DOI:** 10.1101/2020.02.03.931618

**Authors:** Daniel Gyllborg, Christoffer Mattsson Langseth, Xiaoyan Qian, Sergio Marco Salas, Markus M. Hilscher, Ed S. Lein, Mats Nilsson

## Abstract

Visualization of the transcriptome in situ has proven to be a valuable tool in exploring single-cell RNA-sequencing data, providing an additional dimension to investigate spatial cell typing and cell atlases, disease architecture or even data driven discoveries. The field of spatially resolved transcriptomic technologies is emerging as a vital tool to profile gene-expression, continuously pushing current methods to accommodate larger gene panels and larger areas without compromising throughput efficiency. Here, we describe a new version of the in situ sequencing (ISS) method based on padlock probes and rolling circle amplification. Modifications in probe design allows for a new barcoding system via sequence-by-hybridization chemistry for improved spatial detection of RNA transcripts. Due to the amplification of probes, amplicons can be visualized with standard epifluorescence microscopes with high-throughput efficiency and the new sequencing chemistry removes limitations bound by sequence-by-ligation chemistry of ISS. Here we present hybridization-based in situ sequencing (HybISS) that allows for increased flexibility and multiplexing, increased signal-to-noise, all without compromising throughput efficiency of imaging large fields of view. Moreover, the current protocol is demonstrated to work on human brain tissue samples, a source that has proven to be difficult to work with image-based spatial analysis techniques. Overall, HybISS technology works as a target amplification detection method for improved spatial transcriptomic visualization, and importantly, with an ease of implementation.

---

In the era of single-cell transcriptomics, large quantities of data are being produced with various tools and methods. Single-cell RNA-sequencing (scRNA-seq) has revealed the true transcriptomic diversity of cell types within organisms in high detail with such technologies becoming more accessible to laboratories around the world^1–3^. The exponential growth of sequencing large numbers of cells within the last decade has allowed for a systematic assignment to defining cell types, cell states or transitions between them more clear^4–6^. Capturing only static states of cells has been one drawback to sequencing methods and implementation of computational methods trying to overcome this are currently being used, even with *in situ* hybridization data^7–9^. However, one major drawback to scRNA-seq methods that would be difficult to solve computationally is that it requires the cellular dissociation of a tissue to collect cells which could lead to under sampling of vulnerable cell types and more importantly, any spatial context of a cell within a given tissue is lost.

The field of spatial transcriptomics, assigning transcripts a positional location *in situ*, has emerged as a vital tool in the validation of scRNA-seq data and exploring the transcriptomic profiles and cellular architecture across tissues. *In situ* hybridization technologies have come a long way and large consortium efforts are realizing the importance of not only being able to define all the cell types within a human being but also giving them a spatial position in the form of a human cell atlas^10,11^. Image-based transcriptomics have shown the possibility to detect a various range of gene numbers within tissue sections to different degrees of sensitivity and accuracy^12,13^, some used to explore data driven discoveries of biological studies and to view behavior based cellular changes^14–16^.

Spatial distribution of RNA transcripts for cell-type mapping can give further understanding to arrangement of complex tissues^15,17,18^. Multiplexed *in situ* hybridization offers the possibility to explore cellular diversity at subcellular resolution in a upscaled approach. As an example, our lab has developed *in situ* sequencing (ISS) to be used to detect RNA isoforms^19^, transcriptomic distribution^20^, and cell typing across tissue sections^21^. The established ISS method based on barcoded padlock probes (PLPs) and amplification through rolling circle amplification (RCA) has shown robust detection of RNA for various applications, however further improvements are needed to meet the upscaling demands to explore cellular diversity of scRNA-seq data across large tissue areas of various origins.

Here we demonstrate an improved ISS method based on the principles of PLPs and RCA to overcome the inherent limitations of current ISS to address the requirements to investigate spatial mapping of larger gene panels across entire tissue sections. The second iteration of the ISS method uses a sequence-by-hybridization (SBH) chemistry approach to define the location of mRNA molecules within tissue (HybISS: Hybridization-based *In Situ* Sequencing). The unique feature of the ISS method is the probe amplification and the integrated barcoding system that can be decoded across sequential rounds of probing, imaging and stripping. Specifically, the HybISS method takes a new approach to combinatorial barcoding, allowing for more robust detection of molecules and its ability to spatially resolve gene expression *in situ* has been upscaled, supporting the accommodation of larger gene panels, which was one of the main limitations of the sequence-by-ligation (SBL) methods of ISS^20,22^. Here we present HybISS along with an example application to demonstrate the possibility to explore the architecture of human brain tissue and whole mouse brain coronal sections. This results in data sets that can be used to further improve image-based spatial analysis methods such as transcript-spot calling, cell segmentation, and cell typing. Overall, HybISS has benchmarked higher than SBL-based ISS, expanding the possibilities of high-throughput image-based *in situ* transcriptomics.

## Results

### Amplification of padlock probes and sequence-by-hybridization for the detection of RNA transcripts

Similar to previous established ISS methods, PLPs are designed to target specific cDNA sequences from reversed transcribed mRNA *in situ*^20^ (**Figure 1a**). An in-house padlock design pipeline produces multiple gene unique target sequences based on user input parameters (see methods). The customizable backbone of the PLPs contains two parts: a unique ID sequence that is paired to each gene of interest targeted and a general ‘anchor’ sequence shared by all PLPs. Here we design PLPs to target a 30 nucleotide (nt) gene unique cDNA sequence split into two 15 nt arms, and a backbone with a 20 nt ID sequence and a 20 nt anchor sequence, resulting in a final 70 nt long PLP (**Suppl. Figure 1a**). The 20 nt ID sequence is a predetermined sequence unique for the transcript target (e.g. five PLPs targeting five regions of one gene transcript have the same ID sequence). The backbone and ligated target arms of the PLPs are amplified through RCA and results in a rolling circle product (RCP), a sub-micron amplicon detectable with probes and a conventional epifluorescence microscope. The RCP amplicon will contain multiple, repetitive ID sequence targets, allowing for the specificity of complimentary hybridization of readout oligonucleotides termed bridge-probes. Bridge-probes are 39 nt long, 17 bp binding complementary to the ID sequence, a 2 nt noncomplimentary linker, and 20 nt that consists of one of four sequences binding complementary to a 20 bp readout detection probe conjugated to one of four assigned fluorophores (**Figure 1b**, **Suppl. Figure 1b**). Each ID sequence is associated to a combinatorial coding scheme (barcode, e.g. 41233) that is decoded sequentially with a cycle dependent bridge-probe library (**Suppl. Figure 1c**).

**Figure 1:**
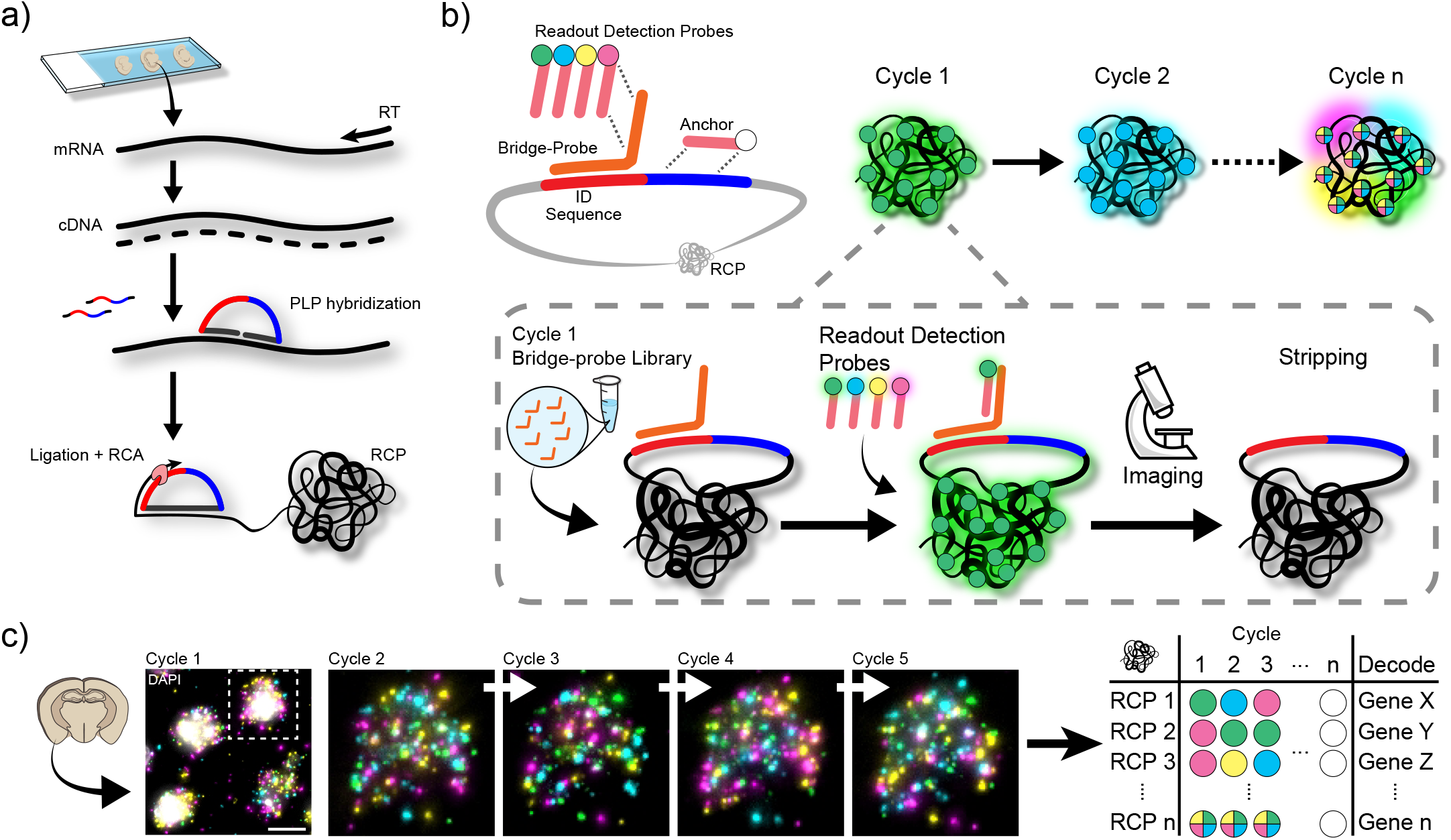
HybISS method overview, hybridization-based in situ sequencing. **a)** Overview of ISS, first reverse transcribing mRNA transcripts to cDNA. Gene specific PLPs target cDNA with juxtaposed ends next to each other that allow for ligation of PLP. Only transcripts that are ligated are enzymatically amplified by RCA. **b)** Schematic overview of HybISS. Every cycle consists of hybridizing bridge-probes to RCPs and reading them out with fluorophore conjugated readout detection probes. For sequential cycles, bridge-probes are then stripped off to allow for rehybridizing next round of bridge-probes. **c)** Example images of 5 cycles of a single cell. Sequential cycles with different bridge-probe libraries allows for the decoding of target transcripts within a cell. Scale bar: 10 μm.

Ligation of juxtaposed 15 nt PLP target-end arms allows for high specificity and RCA of the targeted transcripts in position. Sequential rounds of bridge-probe hybridization, readout detection probing, imaging and stripping allows for a highly multiplexed assay without the round limitation of previous SBL-based ISS methods (**Figure 1b**). Currently, as presented here, HybISS is set up with four readout detection probes per cycle. This allows for a target panel of genes (*G*) that can be measured in a combinatorial manner with four fluorophores (*F*) per cycle (c), and thus *G*=*F*^*c*^, without any cycle limitation and currently showing excellent tissue integrity and RCP maintenance over ten cycles (**Suppl. Figure 1d, e**), roughly calculated to be a loss of 0.28% RCPs per round. Additionally, unlike SBL-based ISS method which is limited to four fluorophores (one per barcode base: A, T, C or G), the number of fluorophores possible in HybISS is limited only by a microscope’s ability to distinguish fluorophores, allowing for increased combinatorial capabilities with proper dye and filter selection. Furthermore, subset bridge-probe panel selection allows for the flexibility to do successive pools of panels on the same tissue sample in the case of overcrowding probes and does not compromise experimental design for probe selection as individual gene readouts can be excluded by simply removing corresponding bridge-probe and done in later cycles (**Suppl. Figure 1f**). This has not been possible in previous ISS methods in that every PLP would be detected every round and problematic genes such as high expressers would have to be run separately. Decoding is performed through sequential combinatorial decoding scheme similar to previous ISS methods, where each barcode has a hamming distance of at least two, allowing for correction and probabilistic assignment of barcodes with better accuracy (**Figure 1c**)^20,21^. Moreover, with bridge-probes, this now also allows the possibility for further error correction by including ‘zeroes’ in the decoding scheme where bridge-probes are simply excluded for a subset of genes in different rounds of probing. All these new features integrated into HybISS increases its flexibility in design of experiments.

### HybISS compared to SBL-based ISS using reference gene panel for benchmarking

In order to demonstrate some of the improvements of HybISS technology, we compare it to SBL-based ISS by designing sets of PLPs to bind identical target sequences of a panel of mouse brain reference genes (*Actb*/*Gapdh*/*Pgk1*/*Polr2a*) and PLP backbones to support the different SBL and SBH sequencing chemistries, resulting in the same length PLPs (**Suppl. Figure 2a, Suppl. Table 1**). SBL and SBH experiments were run in parallel on sequential mouse brain coronal sections showing even distribution of reference genes across tissue sections (**Figure 2a**). First, maximum signal intensities were measured for 100 RCPs per channel per region of interest (ROI) across three comparable regions (**Suppl. Figure 2b**). Measured signal intensities showed a significant increase for SBH-based HybISS across comparative channels (**Figure 2b**). Additionally, intensity was measured across a 21-pixel line centered at random RCPs within the ROIs to measure relative intensity to background. Not only was there an increase in average intensity values, background noise was also reduced in some channels of HybISS and importantly increased signal-to-noise in all channels measured (**Figure 2c, d**). This indicates a more robust detection method, distinguishing RCPs with more ease due to better signal-to-noise. Comparable results were found in additional cycles (**Suppl. Figure 2c**) and this was controlled for by hybridizing only anchors to the respective RCPs and, as expected, no differences seen in intensity values, suggesting no effect on PLP design and in initial steps from hybridizing PLPs to RCA between the two iterations of ISS (**Suppl. Figure 2d, e**). Overall, HybISS benchmarks higher than SBL ISS in more robust signals

**Figure 2:**
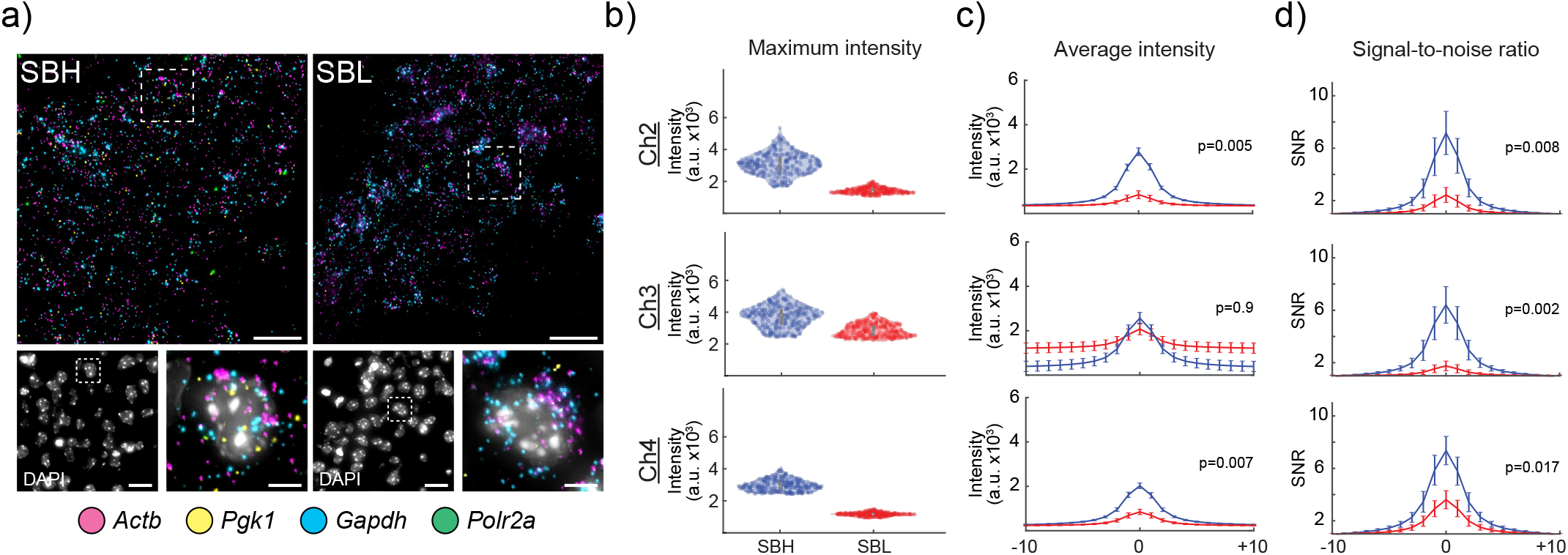
Comparison of HybISS vs SBL-based ISS. **a**) Representative images on the distribution of reference genes in sequential mouse coronal sections targeted by SBH- or SBL-based chemistries. Scale bar: 20 μm, inset 5 μm. **b**) Max intensity of 300 RCPs from three ROIs in each channel measured, comparing SBL- and SBH-based chemistries. Ch2=AF488, Ch3=Cy3, Ch4=Cy5 **c**) Average intensities of measured RCPs in ROIs. Pixel intensity measured across a 21-pixel line bisecting RCPs (red=SBL, blue=SBH). **d**) Outer 2-pixel measurements at each end of 21-pixel line were used to calculate background noise. Each pixel intensity measurement was divided by the noise to display signal-to-noise ratio across the RCPs from (c) (red=SBL, blue=SBH).

### HybISS performed on whole mouse brain coronal section for purpose of gene transcript detection

To demonstrate the potential and validate HybISS, we explored multiplexing in mouse brain sections. The benchmarking gene panel used was curated by the SpaceTx Consortium (part of the Chan Zuckerberg Initiative and Human Cell Atlas project)^10,23^, out of which 119 genes were selected for HybISS PLP design (**Suppl. Table 1**). This probe panel was designed to represent expression of genes to distinguish various cell types within the primary visual cortex^5^ but was applied here to a whole coronal adult mouse brain section (approximately 60 mm^2^, 10 μm thick). Being able to infer data over large areas of the central nervous system, or any tissue, is desired but current spatial methods are limited in their throughput possibilities, imaging only small regions of interest with data storage and time becoming limiting factors. Here we were able to explore tissue architecture in the entire mouse coronal section in an efficient manner, requiring minimal additional input for increased data output. We show the robustness to detect hundreds of target genes and map them in a whole section (**Figure 3a, b**). HybISS was able to resolve the list of genes after combinatorial decoding from five sequential cycles (**Figure 3b, c**), with many genes showing discrete patterns, including laminar structures of cortical tissue (**Figure 3d**). Various genes have very distinct spatial distribution throughout the mouse brain and can be used to validate the method by comparing to other reference atlases such as the Allen Mouse Brain Atlas (2019)^24^ (**Suppl. Figure 3a**). Here we confirm the expression distribution of individual genes not only in the visual cortex but also regions like the hippocampus and thalamus. Although mouse tissue provides a valuable data resource, to understand relevant biological processes in humans, the study of human samples is quintessential.

**Figure 3:**
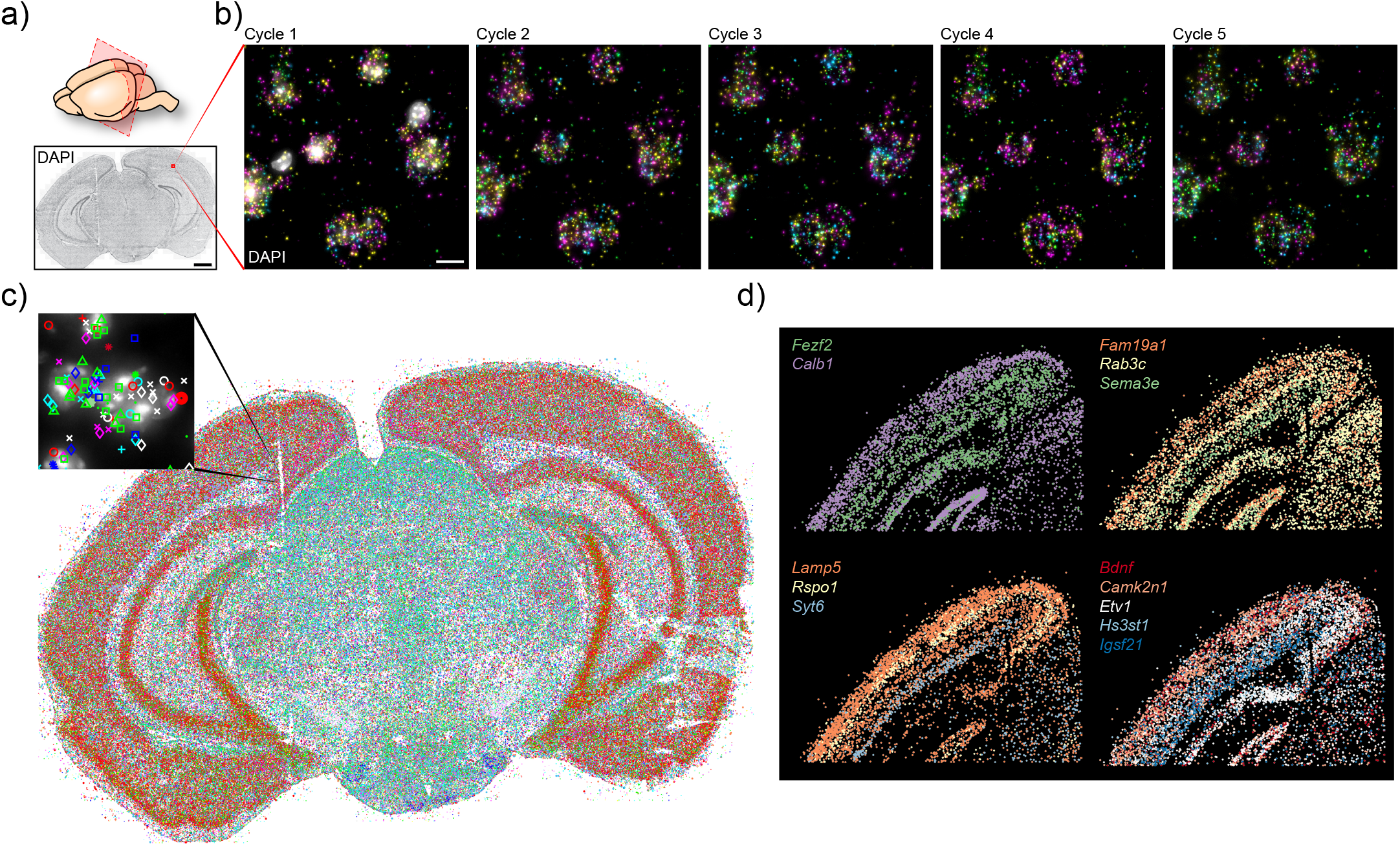
HybISS on mouse coronal brain section. **a**) Whole mouse coronal section used for HybISS. Scale bar: 1 mm. **b**) Representative images of HybISS using PLPs to map 119 genes over 5 cycles in section. Only first image includes counterstain for nuclei with DAPI. Scale bar: 10 μm. **c**) MATLAB output of the decoding of 119 genes across entire section, each color/symbol marking a single transcript detected. Inset shows zoomed in representative image of gene marker plot mapped on DAPI image. **d**) Selection of a subset of genes shown in (c) that have a distinct spatial laminar distribution within the neocortex.

### Multiplexing with HybISS on human brain tissue

Data obtained from human samples would be more relevant and insightful but is limited due to restricted availability of good quality samples and difficulty in adapting and scaling experimental methods. Human tissue regions are much larger than model organisms and therefore investigating comparable regions to mice in humans today still requires significant steps in multiple fields. Well preserved human samples to be used for various single-cell methods are rare and are known to be difficult to work with compared to mouse counterparts where fresh tissue and proven protocols are more abundant. Being able to extract all information possible from a single tissue would provide a wealth of information, and is critical when studying damaged tissue such as the case for neurodegenerative disorders. Furthermore, determining exact anatomical positioning in human samples is more difficult due to their increased area and lack of good reference controls as compared to mouse samples.

Here, HybISS was performed on three sections of human brain tissue from the middle temporal gyrus^25^ (approximately 25-30 mm^2^, 10 μm thick) (**Figure 4a**). One inherent problem with human brain samples is lipid-containing residues of lipofuscin that causes strong autofluorescence in multiple imaging channels and is found in many tissues of aging humans. Many spatial methods and imaging techniques are sensitive to this autofluorescence and overcoming this with any quenching methods does not completely solve the problem and computational clearing strategies are not perfect and require variable user input. Here, we implement HybISS together with simple autofluorescence quenching (TrueBlack Lipofuscin Autofluorescence Quencher, TLAQ) that overcomes this problem in human brain sections and does not require any additional advanced microscopic tools or computational clearing methods (**Figure 4b**). Testing with a human reference gene panel (*ACTB*/*CYC1*/*ACTG1*/*NDUFB4*), in almost all imaged channels without treatment where lipofuscin was present, RCPs could not be distinguished from the background of lipofuscin (**Suppl. Figure 4a, b**). This was further corroborated with investigating crosstalk between channels with and without TLAQ treatment, indicating the effect of lipofuscin on background noise and true signal determination (**Suppl. Figure 4c, d**). TLAQ allows for clear separation of signals in imaged channels from the background and it only adds five minutes to the protocol and a one-time treatment maintains well over imaging cycles.

**Figure 4:**
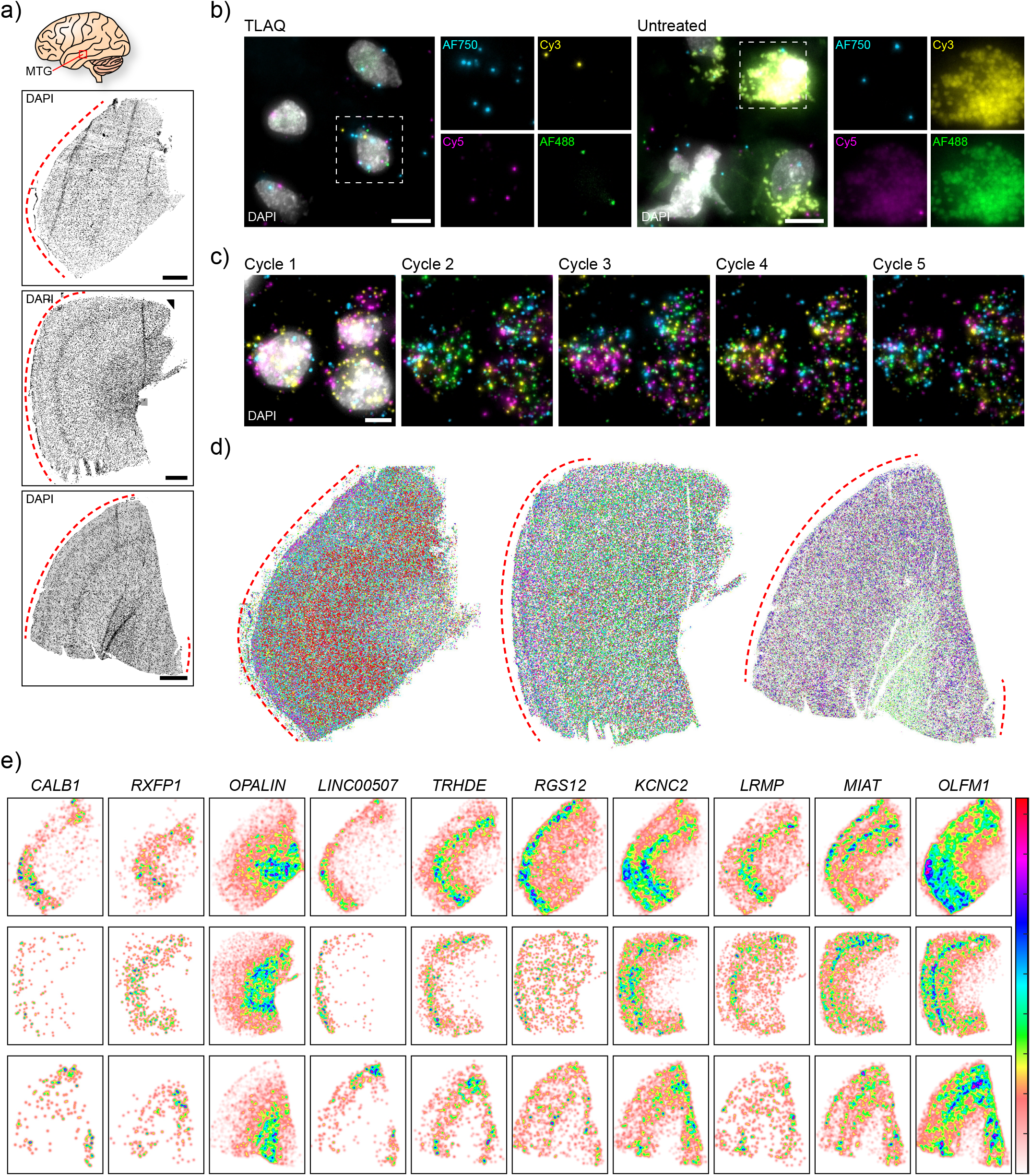
HybISS on human middle temporal gyrus brain sections. **a**) DAPI nuclei stain of human tissue sections from middle temporal gyrus. Dashed red line demarcates the outer surface of tissue section. Scale bar: 1 mm. **b**) Representative images of the effects of lipofuscin in human brain tissue that can be treated with TLAQ and HybISS amplification overcomes any residual background noise. Scale bar: 10 μm. **c**) Magnified field of view from section in (a) of several cells across 5 cycles of HybISS. First cycle includes DAPI to show nuclei location. Scale bar: 5 μm. **d**) Spatial distribution of decoded HybISS transcripts of 120 gene panel across the three tissue sections. (note: markers/colors for genes are not shared between sections) **e**) Kernel density estimation plots for a subset of individual gene transcripts that show distinct spatial distribution, including laminar anatomy of cortical tissue.

With only this additional treatment, we applied HybISS and were able to resolve a similar SpaceTx Consortium curated gene list from human middle temporal gyrus single-nucleus RNA-sequencing data from which a subset of 120 were targeted and decoded (**Figure 4c, d**). We are able to show comparable results to mouse sections without any additional difficulties, indicating a robust method to be used for spatial gene-expression profiling investigations. We further examined the spatial distribution of various genetranscriptswithkerneldensityestimationplotsacross the three sections with many showing distinct patterns, including laminar structures of cortical tissue (**Figure 4e**). Collectively, this data shows a proof of concept that it is possible to do multiplexed *in situ* hybridization studies in human tissue in a high throughput and robust manner.

## Discussion

The HybISS method is a further advancement of the ISS technology in that it strives to meet the requirements and demands in the spatial transcriptomics field without compromising the throughput and area limitation factors that many other techniques are running into. We show the ease of application to human brain tissue with a large gene panel selection to be able distinguish gene transcripts across entire tissue sections, including whole mouse brain coronal sections. The simplicity and flexibility of HybISS makes it a versatile image-based spatial transcriptomic method that can be easily implemented and adapted for a wide variety of scientific questions. Customizing gene lists will be vital as it is a targeted approach and curating custom gene panels from scRNA-seq data, such as probabilistic cell typing by in situ sequencing (pciSeq)^21^, to create cell-type atlases will provide a useful tool in the HybISS method.

Duetotheamplificationmethodof RCA, anepifluorescence microscope using widefield objectives, scanning large regions with good resolution is possible. Amplification-based strategies also aid in overcoming background noise found in many tissue sources. HybISS shows an increase in RCP intensity compared to previous SBL-based methods that allows for robust spot calling, decoding and processing of data. Cleaner and more reliable data aids in downstream analysis such as creating gene networks^26^, cell typing^21^, and molecular maps^27^.

The full potential and implementation of HybISS has not yet been reached. Fine tuning of PLP and target sequence design can increase efficiency, henceforth improve confidence in analysis such as cell typing, although not necessary as even data with lower detection efficiency can yield comprehensive cell maps^21^. Larger gene panels (several fold increase) will be feasible with little to no adjustment to the method presented here. Additionally, as with previous ISS methods, HybISS and PLPs can be used to detect single nucleotide variations due to their high specificity and sensitivity to be able to distinguish point mutations^28^ or editing sites^19^ in tissues for example. SBL-based ISS has already shown its power in other non-neuronal tissue, in the form of cancer diagnostics^20,29^ and tuberculosis granuloma^30^ among others. Therefore, we see no reason for HybISS to improve multiplexed transcript detection in other fields.

We see the great potential of implementing HybISS in spatial cell atlas projects. SBL ISS has been demonstrated to assign cell types in mouse hippocampus^21^ and neural crest development^18^. ISS has also been applied and analyzed together with scRNA-seq data and data from untargeted transcriptome-wide Spatial Transcriptomics^31^ to develop a spatiotemporal atlas of the developing human heart^32^. As demonstrated here on human tissue, the capacity is in place to start creating a comprehensive spatial reference map for the Human Cell Atlas^10^.

Moreover, we provide an image-based spatial transcriptomic data set that can be used to develop computational spatial methods, including Starfish^23,33^, in parallel to further improve the technique. We find the unique insight into the human cortical tissue to be of interest to all and dissemination of such information important. Applying further advancements that have been incorporated, at least in part, to other spatial techniques to HybISS are currently being pursued to try to push the capabilities of the technology. This could be in the form of microfluidic devices to automate production^34^, reducing time, or investigating how these cells are actually arranged in a three-dimensional space and not simply sequential sections.

Implementing tools to further our knowledge of the human body will give us a better understanding of it when in the diseased state. Just like the sequencing of the human genome changed the perspective of human life, a global effort has put us in an era where we are about to define all the cell types within the body. How these cells are organized, connected, and function will give insight to what happens when things go wrong. Here we present HybISS as an *in situ* image-based spatial technology that is reliable, robust, and scalable to meet the emerging demands of the single-cell field.

## Acknowledgements

We thank The Chan Zuckerberg Initiative, an advised fund of Silicon Valley Community Foundation for funding of the SpaceTx consortium. We also thank the Familjen Erling-Persson Foundation, the Knut och Alice Wallenberg Foundation, Swedish Brain Foundation (Hjärnfonden) for generous support of this work. We thank all members of the SpaceTx Consortium for their work contribution that made this possible. We thank Mats Nilsson lab members for their insight and comments.

## Author contributions

D.G. designed and performed the experiments, analyzed data, and wrote the manuscript. C.M.L. performed experiments, analyzed data. X.Q., S.M.S., and M.M.H. analyzed data. E.S.L. and SpaceTx Consortium provided tissue and gene list curation. M.N. conceived and supervised the study. All authors read the manuscript and provided feedback.

## Code and availability

All code is available online at https://github.com/Moldia.

## Supplemental information

Supplemental information contains four figures and four tables.

## Methods

### Protocol

A step-by-step protocol can be found online at protocols.io: dx.doi.org/10.17504/protocols.io.xy4fpyw

### Gene selection

Gene panels were curated both manually and computationally. The panels were based on single-cell RNA-seq data from mouse primary visual cortex^5^ and human middle temporal gyrus^25,35^. Gene list panels were generated as part of the Chan Zuckerberg Initiative SpaceTx Consortium, Working Group 2 and 3. From the curated ranked gene list, a subset was selected to perform HybISS.

### Probe design

Oligonucleotides for PLPs, bridge-probes, detection readout probes, and anchors were ordered from Integrated DNA Technologies (IDT). Stocks of 100 μM (IDTE buffer, pH 8.0) for bridge-probes and detection readout probes were stored at −20°C. PLPs were ordered as large pools and phosphorylated with T4 Polynucleotide Kinase (New England Biolabs). Sequences of all oligonucleotides can be found in **Supplementary Table 1-4**. Target sequences for the selected genes were obtained using in-house Python padlock design software package that utilizes ClustalW and BLAST+ (https://github.com/Moldia/multi_padlock_design) with the following parameters: arm length, 15; Tm, low 65°C, high 75°C; space between targets, 15. After targets sequences were obtained, five targets were selected per gene. If fewer targets were found then only those were chosen. The backbone of the PLPs include a 20 nt ID sequence obtained from the Affymetrix GenFlex Tag 16K Array and a 20 nt sequence ‘anchor’ that is common among subsets of PLPs, a total six of these sequences were used for all PLPs. The decoding scheme for then designing bridge-probes was generated via an in-house MATLAB script that specifies number of positions and minimum base difference between two individual barcodes, hamming distance set to two for five cycles of decoding (https://github.com/Moldia/iss-analysis/tree/master/lib/barcode).

### Tissue collection and preparation

All animal procedures were approved by the Institutional Animal Care and Use Committee at the Allen Institute for Brain Science (Protocol No. 1511). Mouse tissue was obtained from the Allen Brain Institute under the SpaceTx consortium. Fresh whole mouse brain tissue cryopreserved in optimal cutting temperature (OCT) media was stored at −80°C until sectioning. Tissue was sectioned with a cryostat (CryoStar™ NX70) at 10 μm and collected on SuperFrost Plus microscope slides.

Human postmortem tissue collection and processing was performed as previously described in Hodge *et al.*^25^, in accordance with the provisions of the United States Uniform Anatomical Gift Act of 2006 described in the California Health and Safety Code section 7150 (effective 1/1/2008) and other applicable state and federal laws and regulations. Human tissue was received from the Allen Brain Institute under the SpaceTx consortium. 10 μm thick sections were received already mounted on adhesive microscope slides on dry ice and stored at −80°C until used. Tissue from −80°C was first allowed to reach room temperature (RT) (approximately 5 min) before 3% v/v formaldehyde fixation (5 min mouse, 30 min human tissue).

### HybISS

We refer to the accompanying www.protocols. io link for an in-depth description on the method, reagents used and their concentrations. Briefly: *Reverse transcription*: After fixation, tissue sections were permeabilized with 0.1 M HCl for 5 min and washed with PBS. SecureSeal™ Hybridization Chambers (Grace Bio-Labs) were applied around tissue sections and mRNA was reverse transcribed priming with random decamers, RNase inhibitor and reverse transcriptase (BLIRT) overnight at 37°C. *PLP hybridization and ligation*: Tissue sections were fixedfor40minpostreversetranscriptionandsubsequently washed with PBS. Phosphorylated PLPs were hybridized at a final concentration of 10 nM/PLP, and ligated in the same reaction with Tth Ligase (BLIRT) and RNaseH. This was performed at 37°C for 30 min and then moved to 45°C for 1.5 hours. *Rolling circle amplification*: Sections are washed with PBS and RCA is performed with phi29 polymerase (BLIRT) and Exonuclease I (Thermo Scientific). RCA is performed overnight at 30°C. *Autofluorescence quenching*: Human sections were treated with TrueBlack Lipofuscin Autofluorescence Quencher (TLAQ) (Biotium) per manufactures instructions, for 45 seconds and immediately washed with PBS. SecureSeal chambers removed. *Bridge-probe hybridization*: Bridge-probes (10 nM) were hybridized at RT for 1 hour in hybridization buffer (2XSSC, 20% formamide). *Readout detection probe hybridization*: This is followed by hybridization of readout detection probes (100 nM) and DAPI (Biotium) in hybridization buffer for 2 hours at RT. Sections are washed with PBS and sections are mounted with SlowFade Gold Antifade Mountant (Thermo Fisher).

### Imaging

Imaging was performed using a standard epifluorescence microscope (Zeiss Axio Imager.Z2) connected to external LED source (Lumencor^®^ SPECTRA X light engine). Light engine is setup with filter paddles (395/25, 438/29, 470/24, 555/28, 635/22, 730/40). Images obtained with a sCMOS camera (2048 × 2048, 16-bit, ORCA-Flash4.0 LT Plus, Hamamatsu), automatic multi-slide stage (PILine, M-686K011), and Zeiss Plan-Apochromat objectives 20x (0.8 NA, air, 420650-9901), 40x (1.4 NA, oil, 420762-9900). Filter cubes for wavelength separation included quad band Chroma 89402 (DAPI, Cy3, Cy5), quad band Chroma 89403 (Atto425, TexasRed, AlexaFluor750), and single band Zeiss 38HE (AlexaFluor488). In SBL-based ISS, one of the base libraries is either conjugated to TexasRed or Atto425 fluorophores that requires substitution of the 555/28 filter paddle with a compatible 575 one to image TexasRed.

### Stripping

After imaging, sections are prepared for sequential cycles. Readout detection probes and bridge-probes are stripped off RCPs. This is performed by 5 washes with 2XSSC and then sections are incubated in stripping solution (65% formamide, 2XSSC) for 30 min at 30°C. This is followed by 5 washes with 2XSSC. Now the next cycle of bridge-probes can be hybridized as previously.

### Image analysis

Imaging data was analyzed with in-house custom software that handles image processing and gene calling. All code is written in MATLAB and is freely available at (https://github.com/Moldia). Many steps follow previous publications ^20,21,29,30^.

Multispectral imaging for channels occurs for multiple cycles. Each image consists of multiple tiles that contain a 10% overlap to allow for stitching alignment between fields of view. A z-stack focal depth of 10 μm with a spacing of 0.5 μm deep allows for coverage of entire tissue. Tiles were stitched together using Zeiss ZEN software and then maximum intensity projection was performed in Zeiss ZEN software to obtain a flattened two-dimensional image for each channel imaged. After the images had been exported to .tif format, the images were split into multiple smaller images, henceforth referred to as tiles. The tiled images were subsequently top-hat filtered, the RCPs were segmented and the intensity of each RCP was measured, for each channel. Prior to measuring the intensities, there is an alignment step, meaning that RCPs are aligned between cycles. The above steps, after the tiling, were done in CellProfiler 2.2.0. The intensity measurements were saved in a comma separated values and subsequently used to decode the RCPs in MATLAB (repository: https://github.com/Moldia/Tools). These intensity measurements are used to calculate a quality score, defined as follows: the maximum intensity in the set of channels is divided by the sum of the all of the channels. This means that we can have a quality score of 0.25 to 1. Similar intensities in all channels would result in a quality score on the lower end of this spectrum and at the upper end, the opposite, i.e. more dissimilar intensities between the maximum intensity channel and the other channels.

### SBH vs SBL analysis

Each method was performed in parallel on two sequential full mouse brain coronal sections using four PLPs per target gene *Actb*, *Pgk1*, *Gapdh*, and *Polr2a*, allowing for the possibility to decode in one cycle. The protocol for the steps to decode SBL chemistry was as previously published^21^. From the entire imaged sections, for comparison of RCPs quality, and to reduce the computational demands, the sections were divided into subsets. Three regions of interest (ROI) were drawn to cover isocortex (ROI1), hippocampal formation (ROI2) and thalamus (ROI3) (**Supp. Figure 2b**). The size of each ROI was 6000 x 6000 pixels. These new images were then exported as TIFF files. *Intensity measurements over RCPs:* For each channel image, the RCPs were located and the RCPs intensity measured. The intensity was measured over the RCP, 10 pixels in each direction from the middle, which generated a total of 21 intensity measurements. *Average intensity over the RCPs:* The average intensity and the standard deviation over the RCPs was calculated. This was then visualized using line plots. An unpaired t-test was then used to assess the degree of statistical significance, which was assessed by taking the middle five values. Figure 2c: DO3-SBH, n=1216; DO3-SBL, n=2490; DO4-SBH, n=1576; DO4-SBL, n=2760; DO2-SBH, n=417; DO2-SBL, n=1247. Supp. Figure 2c: DO3-SBH, n=1251; DO3-SBL, n=814; DO4-SBH, n=466; DO4-SBL, n=1036; DO2-SBH, n=1058; DO2-SBL, n=2070. *Maximum intensity:* The top 100 intensities were then extracted from the middle of each RCP that was located, generating 100 intensities values from each channel and then done for the three ROIs. The maximum intensities were then visualized with violin plots. *Signal-to-noise ratio:* For each of the RCPs found, the noise was defined as the four outer positions, i.e. pixel number −10, −9, 9 and 10. The mean for these positions was calculated and defined as the noise. The intensity value for each position was then divided by the noise. The mean and the standard deviation for these values were then calculated and plotted. *Control experiment:* In the control experiment (**Supp. Figure 2a**), an anchor probe was hybridized to both sections and the average intensity, SNR and the SNR in the center of RCP was compared between the two chemistries.

### Crosstalk measurements

Crosstalk between channels was assessed by cytoflourograms. For each pixel position, the intensity of one channel is plotted on the x-axis and the intensity of another channel is plotted on the y-axis. This means that the pure signals will populate the paraxial regions and the brightest spots will populate the ends of the axes. Mixed pixels will be in the diagonal area of the plot.

