## Supplemental Figures 1-4 for "Hybridization-based In Situ Sequencing (HybISS): spatial transcriptomic detection in human and mouse brain tissue"

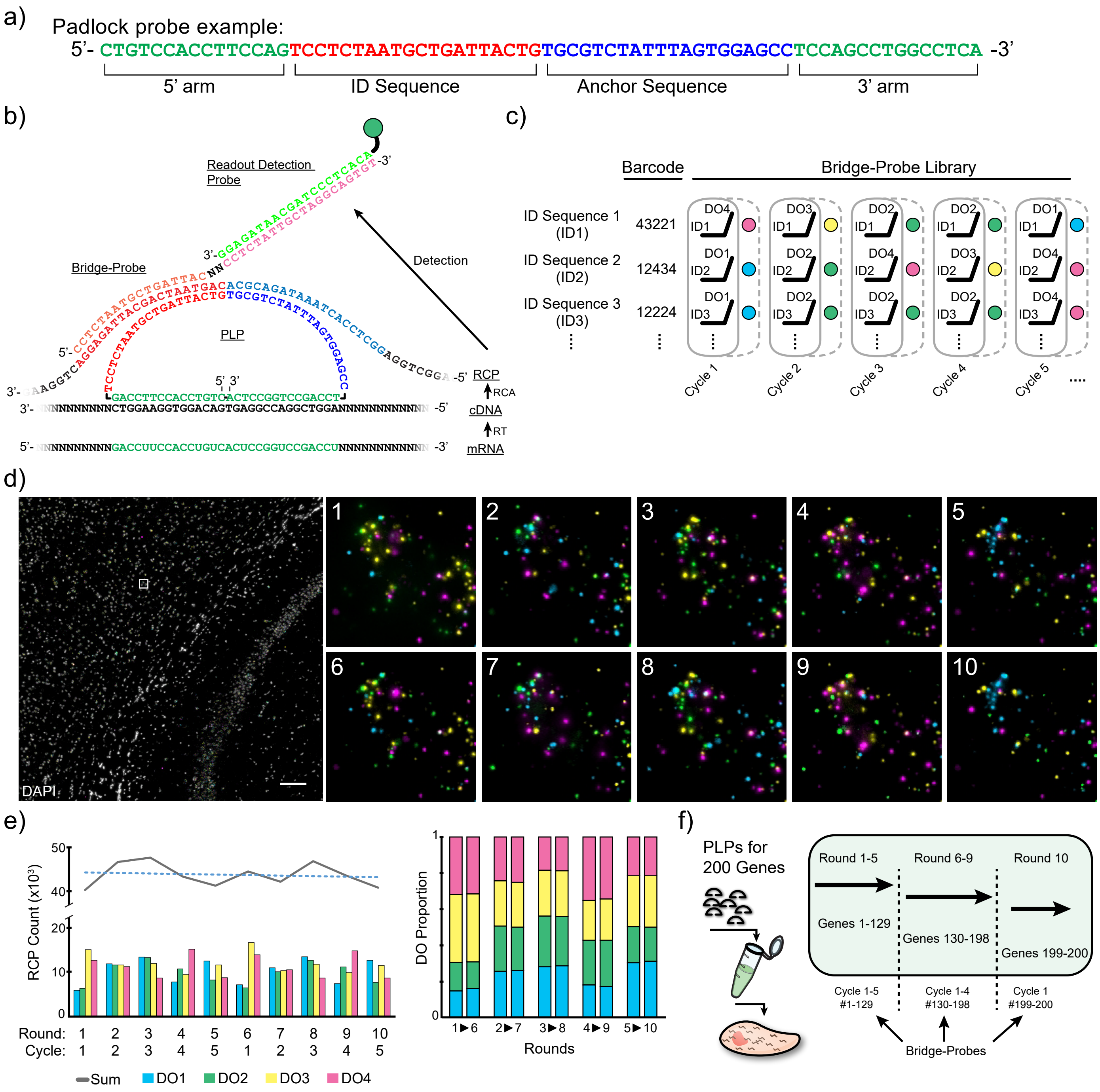


***Supplemental Figure 1: Padlock probe design and multi-round imaging.*** ***a****) Example of a PLP sequence, 5' to 3'. Target recognition sequence (green) is split to each arm allowing for end to end juxtaposed positioning and ligation. Backbone consists of a gene unique 20 bp ID sequence (red) and common 20 bp anchor sequence (blue).* ***b****) Overview of how bridge-probes and readout detection probes are designed considering the 5' to 3' direction of various products from the different steps in the method.* ***c****) Overview example of how bridge-probe libraries function across several cycles that allows for combinatorial decoding.* ***d****) HybISS and multi-round probing. Image used for RCP counting in panel (e). Inset image shows detail across 10 rounds of probing and imaging, 119 mouse gene panel used. Scale bar: 100 µm.* ***e****) RCP counts in ROI (panel (d), 1.4 mm x 1.4 mm) in a 10-round experiment (trendline: y=-125.03x+44630). 5 cycle experiment was repeated on same tissue for round 6-10, giving similar proportions to detected RCPs in comparative cycles (e.g. round 1 vs. round 6).* ***f****) Schematic overview of how RCP readout can be performed in the case of overcrowding by splitting up bridge-probe panels for RCP detection.*

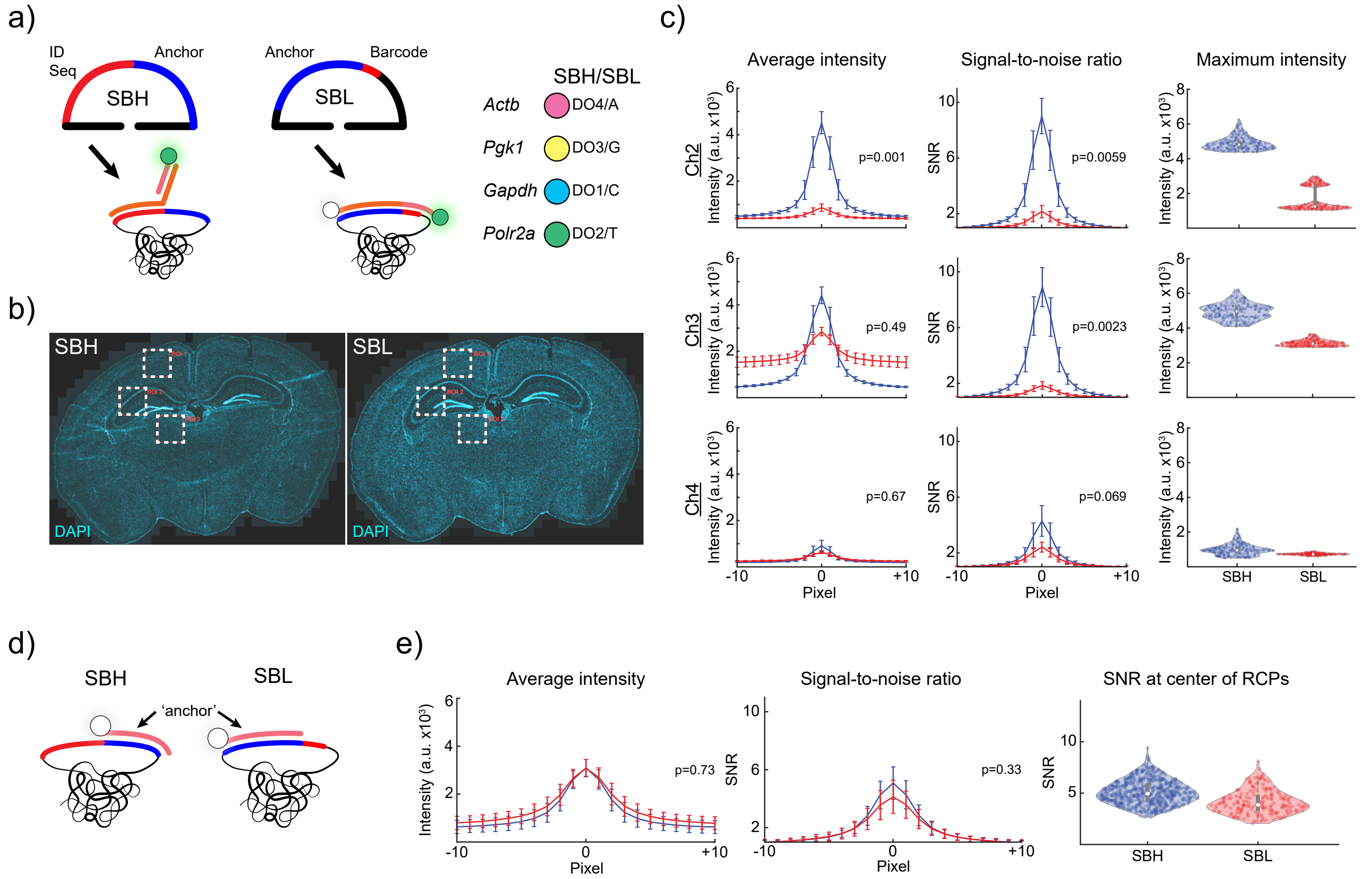


***Supplemental Figure 2: Benchmarking HybISS against SBL-based ISS. a****) Schematic overview of PLP design of SBL- vs SBH-based chemistries.* ***b****) Three ROIs used for analysis in measurement of intensities from sequential sections of mouse brain tissue. DAPI staining for nuclei.* ***c****) Additional measurements for average intensity, and SNR maximum intensity, for a different cycle compared to Figure 2. DO2=AF488, DO3=Cy3, DO4=Cy5.* ***d****) Control experiment schematic to show no difference in intensities when hybridizing anchor (Fluorophore AF750) only between the SBL- and SBH-based chemistries due to both anchors being hybridization only.* ***e****) Measured average intensity across RCPs and the signal-to-noise ratio of those intensities in the control experiment.*

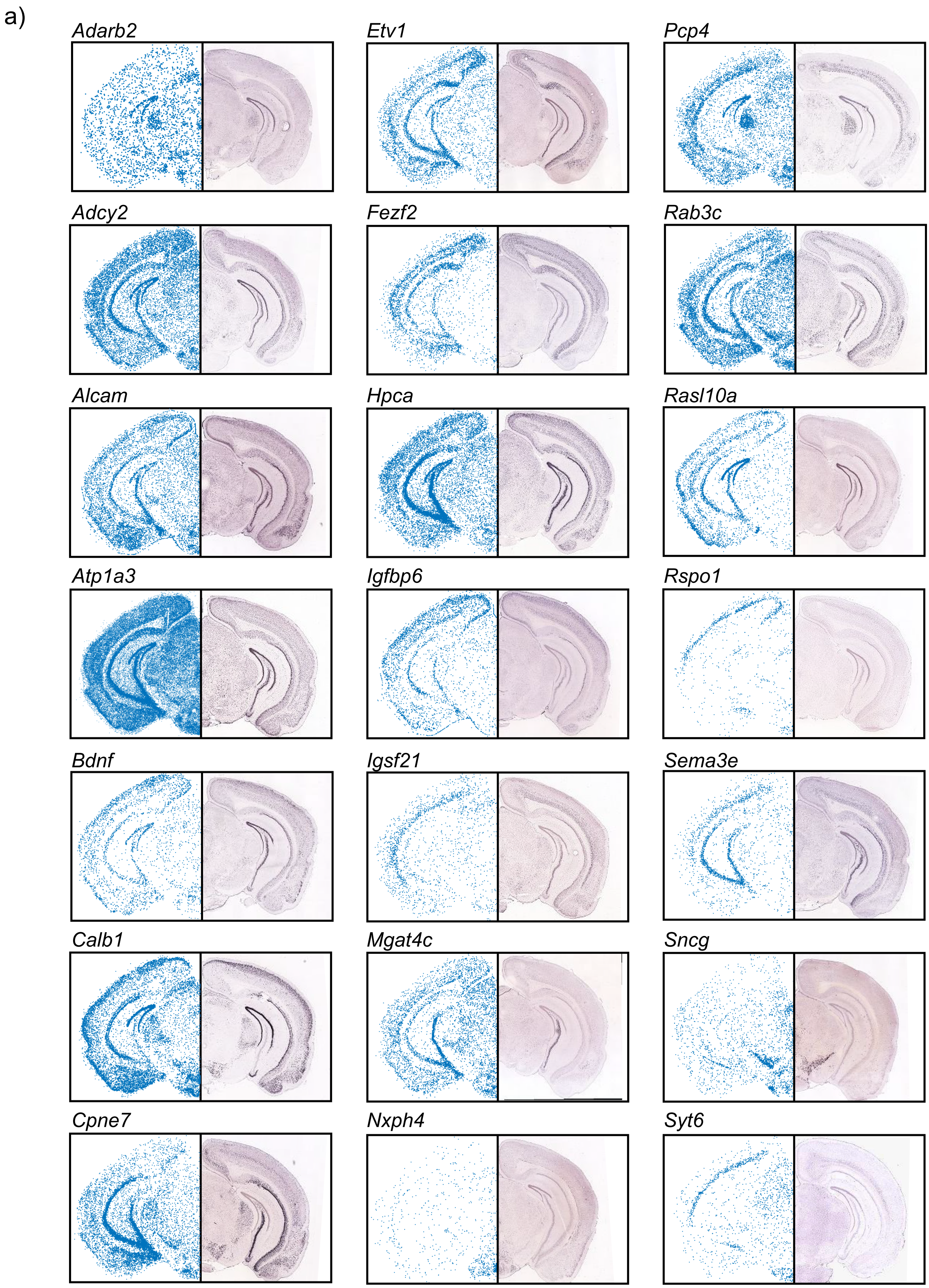


***Supplemental Figure 3: Validation of spatial distribution of mouse gene. a)*** *Spatial distribution of subset selection genes from HybISS method (left, blue) from data in Figure 3 compared to RNA in situ hybridization the Allen Mouse Brain Atlas[24] (right). Image credit: Allen Institute.*

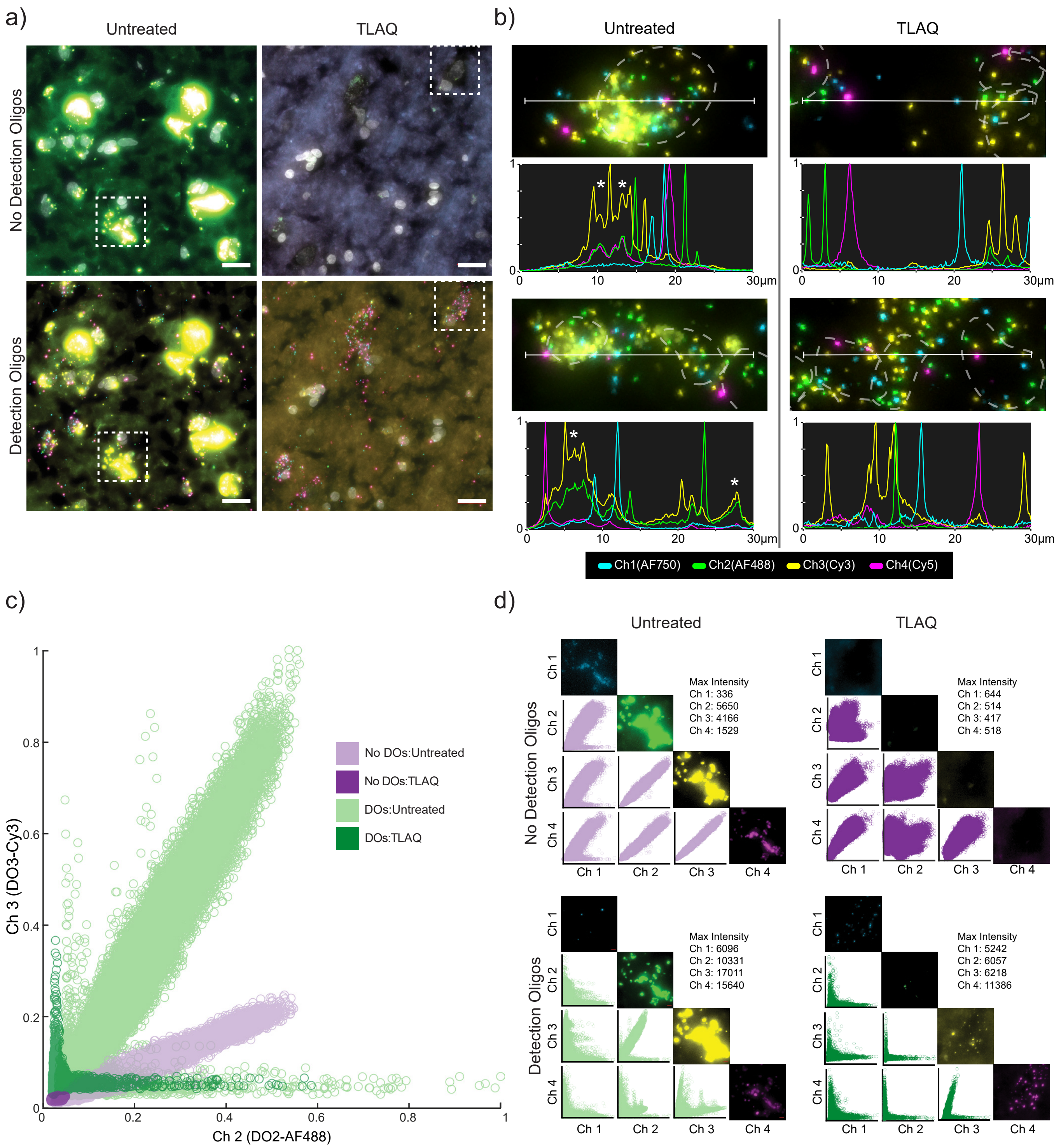


***Supplemental Figure 4: Lipofuscin analysis in human brain tissue. a**) ROIs from human sample experiments used for analysis in panel (c) and (d). A panel of human reference genes (ACTB/CYC1/ACTG1/NDUFB4) were used for probing. Untreated and TLAQ treated samples run in parallel and imaged (top panels) then bridge/ DOs hybridized and imaged again (bottom panels). Outlined inset in each image is shown in panel (d) as an example across different channels. ROI area =1000x1000 pixels, (26569 μm$^2$) Scale bar: 20 μm. **b**) Examples of profile intensities across RCPs, lipofuscin and background noise in untreated and TLAQ treated conditions in human brain sections. Profile intensities are normalized to maximum and minimum of each channel. Dashed line is nuclei outline. Asterisk indicates lipofuscin crosstalk. Profile measured across 30 μm (white line). **c**) Crosstalk between Ch 2 (AF488) and Ch 3 (Cy3) from images in panel (a). Each data point represents a single pixel intensity from 2 channels. All conditions were normalized to maximum intensity value in one of the 4 images (DOs:Untreated). No DOs:TLAQ represents only background noise intensities. **d**) Crosstalk between individual channels for whole image in panel (a). Each plot is normalized to the maximum intensity of the plotted channel in respective image, these max values are displayed in each sub-panel. Center plot in each sub-panel arrangement (Ch 2 vs Ch 3) is same as represented in panel (c) except normalized to different max pixel intensities. Ch 1=AF750, Ch 2=AF488, Ch 3=Cy3, Ch 4=Cy5.*
